# Alteration of brain dynamics during natural dual-task walking

**DOI:** 10.1101/2020.02.27.968164

**Authors:** Federica Nenna, Cao-Tri Do, Janna Protzak, Klaus Gramann

## Abstract

While walking in our natural environment, we continuously solve additional cognitive tasks. This increases the demand of resources needed for both the cognitive and motor systems, resulting in Cognitive-Motor Interference (CMI). While it is well known that a performance decrease in one or both tasks can be observed, little is known about human brain dynamics underlying CMI during dual-task walking. Moreover, a large portion of previous investigations on CMI took place in static settings, emphasizing the experimental rigor but overshadowing the ecological validity. To address these problems, we developed a dual-task walking scenario in virtual reality (VR) combined with Mobile Brain/Body Imaging (MoBI). We aimed at investigating how brain dynamics are modulated during natural overground walking while simultaneously performing a visual discrimination task in an ecologically valid scenario. Even though the visual task did not affect performance while walking, a P3 amplitude reduction along with changes in power spectral densities (PSDs) during dual-task walking were observed. Replicating previous results, this reflects the impact of walking on the parallel processing of visual stimuli, even when the cognitive task is particularly easy. This standardized and easy to modify VR-paradigm helps to systematically study CMI, allowing researchers to control the complexity of different tasks and sensory modalities. Future investigations implementing an improved virtual design with more challenging cognitive and motor tasks will have to investigate the roles of both cognition and motion, allowing for a better understanding of the functional architecture of attention reallocation between cognitive and motor systems during active behavior.

## INTRODUCTION

In our daily activities, we need to walk and interact with environmental cues in order to meet everyday goals. This entails the processing of both external and internal sensory information that help maintaining action goals, react to changing environmental features, and readapt motor programs anytime unexpected events occur. Therefore, despite usually perceived as undemanding, walking involves both sensory and cognitive systems (Hausdorff et al., 2008; Woollacott & Shumway-Cook, 2002). In these highly dynamic situations, our limited attentional resources have to be distributed between the motor and cognitive tasks, potentially causing a cognitive-motor interference (CMI) (for a review, Al-Yahya et al., 2011). This phenomenon has been widely investigated through dual-task walking paradigms and, more recently, through new mobile neurophysiological approaches. Particularly, Mobile Brain/Body Imaging (MoBI; Makeig et al., 2009; Gramann et al., 2011, 2014; Jungnickel et al., 2019) has been proposed as a method for recording brain data during active movement, allowing to gain physiological measurements of the whole brain-body system. The general feasibility of the MoBI concept has been demonstrated and applied to dual-task walking scenarios (Gramann et al., 2010; De Sanctis et al. 2012, 2014; Debener et al., 2012; Duvinage et al., 2013; Hoellinger et al., 2013; Castermans et al., 2014; De Vos et al., 2014; Reis et al., 2014). In a pioneering work, Gramann and colleagues (2010) analysed the brain dynamics of participants during standing, slow and fast treadmill walking, while attending to a visual oddball task. They demonstrated that the oddball P3 known from traditional desktop scenarios can be replicated in paradigms allowing active walking (Gramann et al., 2010).

So far, several cognitive tasks have been used to study CMI during walking, such as visuo-motor reaction time tasks (Patel et al., 2014), word list generation task (Patel et al., 2014), serial subtraction task (Marcar et al., 2014; Patel et al., 2014), stroop task (Patel et al., 2014), texting (Plummer et al., 2015), Go/NoGo (De Sanctis et al., 2014; Malcolm et al., 2015; Beurskens et al., 2016) or oddball task (Gramann et al., 2010; Gwin et al., 2010; Debener et al., 2012; Ladouce et al., 2019; Reiser et al., 2019). These tasks utilized different sensory modalities for the cognitive task including the visual and auditory domain as well as different postural tasks, including sitting, standing, walking on a treadmill or, in a few recent cases, walking over-ground in real contexts (Debener et al., 2012; Plummer et al., 2015; Pizzamiglio et al., 2017; Ladouce et al., 2019; Reiser et al., 2019).

### 1.1 Performance perturbation under CMI

Irrespective of the cognitive task modality, during CMI, a performance deterioration is usually observed in the cognitive and/or in the motor task (Leone et al., 2017). Focusing on the motor behavior, studies on dual-task walking demonstrated alterations in gait velocity, stride length and stride time (Beurskens et al., 2016; De Sanctis et al., 2014; Malcolm et al., 2015; Patel et al., 2014; Pizzamiglio et al., 2017; Plummer et al., 2015). This suggests that gait parameters vary with attentional resources even in healthy adults with good locomotor and cognitive functions. From a cognitive perspective, behavioral costs have been observed in response to both visual (Patel et al., 2014; Plummer et al., 2015) and auditory tasks (Beurskens et al., 2016; Reiser et al., 2019). However, in a direct comparison between reporting the content of a message presented visually and aurally while detecting obstacles, only the presentation of visual but not auditory messages yielded significant differences in task performance (Silva et al., 2019). This suggests that visual tasks, compared to auditory tasks, might generate a more pronounced conflict in resource allocation when concurrently walking because gait also requires visual attention to control the upcoming path (Imai et al., 2001; Nomura et al., 2005; Marigold & Patla, 2008). Focusing on visual tasks, increased visuo-motor reaction times and less correct responses have been observed in the execution of the stroop task when walking compared to sitting (Patel et al., 2014). On the same line, in comparison to the standing condition, a drop in performance under dual-task occurred when walking and concurrently texting (Plummer et al., 2015) or performing a visual oddball task (Beurskens et al., 2016).

### 1.2 Electrophysiological modulations under CMI

Studies on event related potentials (ERPs) investigating CMI mainly focused on the investigation of the P3 component, which is sensitive towards the amount of attentional resources engaged to solve a task (Polich, 2007; Israel et al., 1980a, 1980b). In the frequency domain, several studies showed power spectral density (PSD) parameters to be related to cognitive and/or motor load in stationary dual-task-based studies but not yet in mobile investigations. For the latter, only a few and sometimes contradictory observations have been reported (Gwin et al., 2010; Presacco et al., 2011; Beurskens et al., 2013; 2016; Marcar et al., 2014; Pizzamiglio et al., 2017; Peterson & Ferris, 2018).

#### ERPs of CMI during walking

A decreased **P3 amplitude** at centro-parietal electrodes from single to dual-tasks was often related to the restricted availability of cognitive resources (Polich, 2007). P3 amplitude reductions in mobile EEG walking studies were interpreted as a reflection of a greater demand for attentional resources when walking (Debener et al., 2012; De Sanctis et al., 2014; Malcolm et al., 2015; Ladouce et al., 2019; Reiser et al., 2019). In contrast, a significant increase of P3 amplitude was reported over fronto-central regions when walking as compared to sitting in young participants (De Sanctis et al., 2014). Interestingly, in some cases the P3 amplitude modulation was observed in dual-task walking paradigms even in the absence of behavioral evidence of performance cost (De Sanctis et al., 2014; Malcolm et al., 2015).

**P3 latency** modulations were also observed in dual-task walking scenarios. An earlier P3 onset was observed over frontal and centro-parietal regions when walking compared to sitting in young healthy adults (De Sanctis et al., 2014; Malcolm et al., 2015).

#### PSDs of CMI during walking

Stationary dual-task studies have demonstrated that a tonic increase in theta power (4-7 Hz) together with a decrease in alpha power (8-12 Hz) are often observed over wide areas of the scalp under higher task demand (Klimesh et al., 1999). Focusing on MoBI studies, despite the diversity of task designs and modalities, higher PSD in the **theta** band has been observed when walking compared to standing or resting still (Presacco et al., 2011; Peterson & Ferris, 2018). At the same time, lower **alpha** activity over widespread areas of the scalp (from frontal to parietal cortex, and in some cases over the whole scalp) has also been observed when walking as compared to standing or resting (Beurskens et al., 2016; Peterson & Ferris, 2018). In addition, a decreased alpha power was observed when performing cognitively engaging tasks such as walking in an interactive virtual environment (Wagner et al., 2014) and closed-loop brain-computer interface (BCI) control of a virtual avatar walking (Luu et al., 2017).

For the **beta**-band (12-30 Hz), different results have been observed for dual-task modulations during walking. In established stationary setups, increased beta power has been observed over parieto-occipital brain areas when cognitive load increased (Belyavin & Wright, 1987; Gola et al., 2013) or when increasing the effort to stay alert (Boksem et al., 2005; Lafrance et al., 2000). Further, beta power increases were reported for performing demanding motor tasks (i.e. grasping tasks; Zaepffel et al., 2016) or when walking as compared to resting (Presacco et al., 2011). Focusing on mobile setups, Beurskens et al., (2016) detected a beta power increase over frontal regions when walking while performing a motor task whereby all the effort was attributable to the motor system. This beta power increase was more pronounced for a secondary motor task condition than an additional secondary cognitive task (Go/NoGo task). The same study also showed decreased beta power over frontal and central electrodes when walking while performing the Go/NoGo task with respect to single-task walking. A similar decrease in beta band power was demonstrated at frontal, central and parietal electrodes during single-task walking as compared to standing still (Pizzamiglio et al., 2017). The role of beta band activity in attention processes is well established for numerous thalamic and cortical centers of the visual system (Wróbel, 2000 for a review). However, mechanisms of beta power modulation by cognitive and/or motor task load are still unclear.

**Gamma** power (> 30 Hz) is implicated in active cortical processing and was shown to increase with greater postural instability (Slobounov et al., 2009) which is likely to occur in dynamic situations like walking. Gait cycle dependent modulations in the gamma band have been reported over frontal, central, and parietal cortex of healthy walking adults (Gwin et al., 2011). Marcar et al. (2014) observed an increase of gamma power when walking whilst performing a serial subtraction task. Recently, an increased power at high frequencies was also reported by Peterson and Ferris (2018) when walking as compared to standing. However, frequencies above 30 Hz may be compromised by cranial muscle activity (both facial and neck muscles) (O’Donnell et al., 1974; Goncharova et al., 2003; Whitham et al., 2008). Therefore, neural activity in the gamma frequency band has to be interpreted with caution and relies on analyses tools addressing mixtures of different physiological signals at the channel level.

### 1.3 The importance of ecological validity

So far, a large portion of neurophysiological CMI investigations took place in static and artificial settings. Newer mobile EEG and MoBI approaches assessed CMI with treadmill setups, but only rarely during natural overground walking (Debener et al., 2012; Pizzamiglio et al., 2017; Ladouce et al., 2019; Reiser et al., 2019). This leads to a lack of ecological validity, which plays a prominent role when studying real-world neural dynamics and behaviors (Gramann et al., 2011; 2014). Particularly when investigating dual-task walking using secondary tasks that tax the visual modality, it is necessary to replace static desktop displays and treadmill setups with ecologically more valid but controlled paradigms. This is because orienting movements of our head naturally occur during everyday additional visual tasks and gait parameters might be dynamically adapted dependent on the task at hand. This is not possible during treadmill walking where participants have to keep up a certain gait velocity to secure their position on the treadmill while focusing on a visual display directly in front of them.

Visual information processing while walking is important for at least two reasons. First, walking requires visual control of the upcoming area to avoid falls or collisions (Imai et al., 2001; Nomura et al., 2005); as a consequence, an additional visual task will rely on the same resources which are necessary to visually control walking and thus compete (Wickens et al., 1983; Wickens, 2002). Secondly, the sensory features and position of additional visual stimuli (eccentricity in the visual field) have a strong impact on visual information processing (Carrasco et al., 1995; Staugaard et al., 2016). Therefore, detecting peripheral and potentially less salient visual stimuli might require more attentional resources compared to more centrally ones, revealing a higher dual-task cost when concurrently walking.

In the present study, we simulated an everyday situation through virtual reality (VR) in which people stood or walked freely while discriminating and responding to external visual stimuli. We hypothesize a performance decrease in the visual discrimination task, particularly for more peripherally as compared to more centrally presented visual stimuli, when the cognitive and motor tasks had to be performed simultaneously. Higher perceived subjective mental load was also expected when walking under dual-task. As reported in previous studies (De Sanctis et al., 2014; Malcolm et al., 2015), we expected to find the P3 component evoked by the onset of a visual stimulus to peak earlier over frontal and central areas of the brain, and to show lower amplitude over centro-parietal areas in the dual-task condition as a consequence of the reallocation of cognitive resources under dual-task conditions. Finally, we also expected to observe higher theta and lower alpha power from frontal to parietal brain areas as a function of the mental load evoked by the dual-task. Moreover, increased beta and gamma power were hypothesized under dual-task walking compared to the single-task situation.

## MATERIALS AND METHODS

### 2.1 Participants

25 right-handed participants with normal or corrected to normal vision as well as normal color vision were recruited. All participants reported to be in good health and free of any neurological impairments. They also reported the absence of medication containing psycho/neuroleptics, as well as intoxicant use within the last 24 hours prior to the experiment. 3 participants were excluded from the analysis due to technical issues. The remaining sample included data from 22 participants: 6 females (age range: 20-31 years, M = 25.5, SD = 3.92) and 16 males (age range: 21-34 years, M = 27.2, SD = 4.48). Before the experiment, participants were asked to report their height in centimeters for adapting the virtual environment (height range for female: 170-172 cm, M = 170.33 cm, SD = 0.83 cm; height range for male: 170-181 cm, M = 174.38 cm, SD = 4.15 cm). The study was approved by the TU Berlin ethics committee. All participants gave written informed consent and were recruited through the local online participant portal (https://proband.ipa.tu-berlin.de). Participants obtained credits for compensation.

### 2.2 Technical set-up

To investigate the neural correlates of CMI, we designed the following experimental framework following the MoBI set-up shown in figure 1. It consisted of a 128 channels EEG MOVE system (Brain Products GmbH, Gilching, Deutschland), in combination with actiCAP (Easycap GmbH, Herrsching, Germany), and a VR headset (ACER WMR; 2.89”, 2880 x 1440 resolution, refresh rate of 90 Hz, 100° field of view with a weight of 440 grams). The headset was tethered to a Zotac gaming computer (Zotac PC, Intel 7th Gen Kaby Lake processor, GeForce GTX 1060 graphics, 32GB DDR4-2400 memory support, Windows 10 OS) placed in a backpack. The Zotac system was extended with two batteries that allowed swapping them approximately after three blocks of experimental session (circa 40 minutes) without shutting down the VR. Participants were projected into a virtual environment (VE) designed in Unity (2017.3). In addition, a prototype of VR-EEG adapters (Wenzel, 2018) was used to reduce the mechanical pressure on frontal and occipital channels of the EEG cap induced by the VR goggles. The adapters further minimized the signal to noise ratio by reducing headset movements accompanying locomotion. Both behavioral and neural data streams were synchronized via Lab Streaming Layer (Kothe, 2014).

**Fig. 1:**
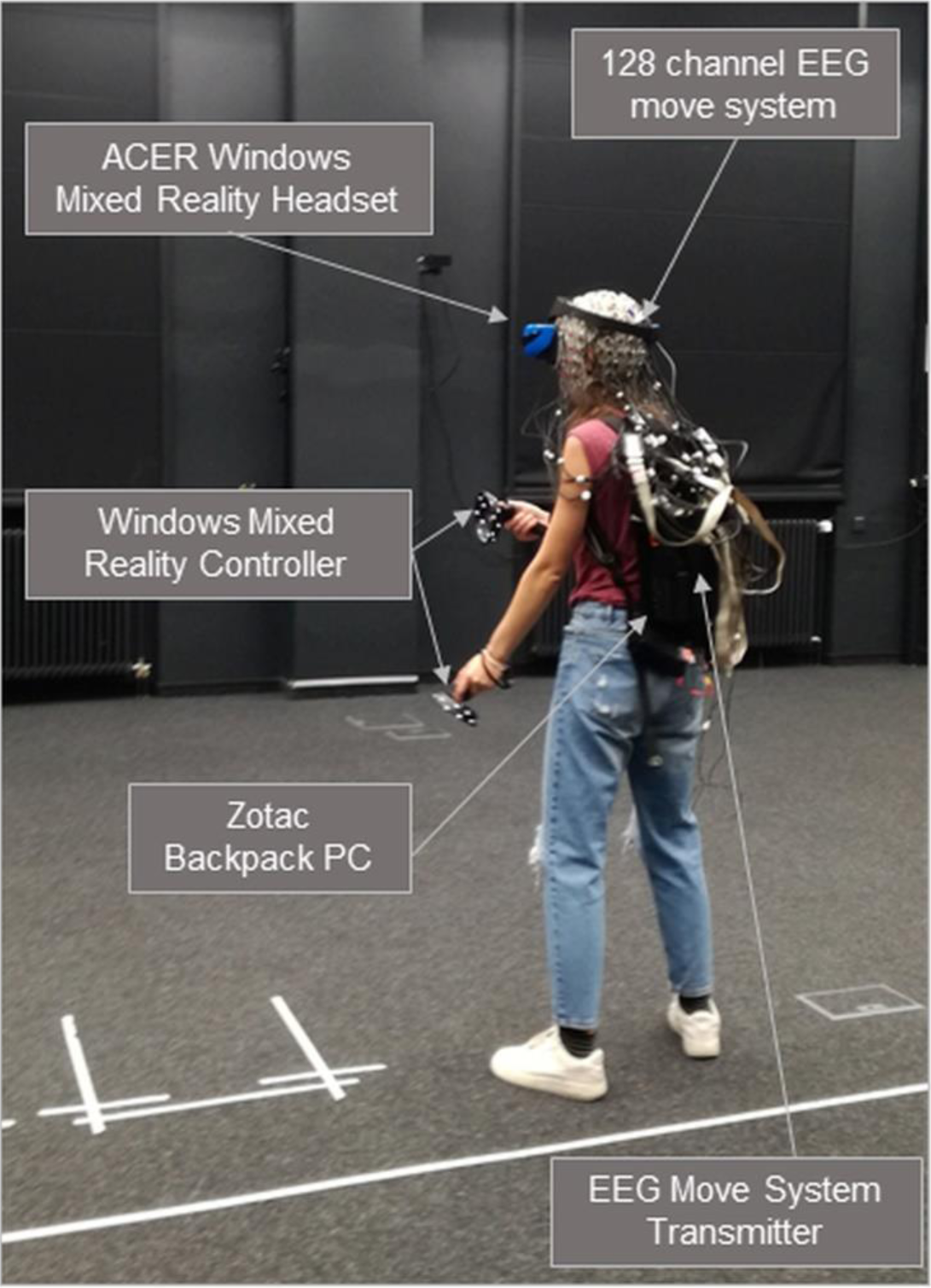
Overview of the technical setup for the MoBI experiment.

### 2.3 Experimental design

The experiment took place at the Berlin Mobile Brain/Body Imaging Laboratories (BeMoBIL), with a dedicated room providing an experimental space of 150 m^2^ for participants to move around without restrictions. The virtual space provided an elliptical path that was 10.8 m long and 2.5 m wide. Data collection took place in one single experimental session. During the training phase, participants were asked to walk along the oval path and to follow a red moving sphere (figure 2). Prior to the experiment, each participant was able to adjust the speed of the sphere to their preferred natural walking pace using the controller in a training session. The speed was then kept constant during the experiment. In the second part of the training phase, participants were instructed to follow the sphere while performing a visual discrimination task. This part consisted of 15 trials with the aim of familiarizing with the task and the virtual environment.

**Fig. 2:**
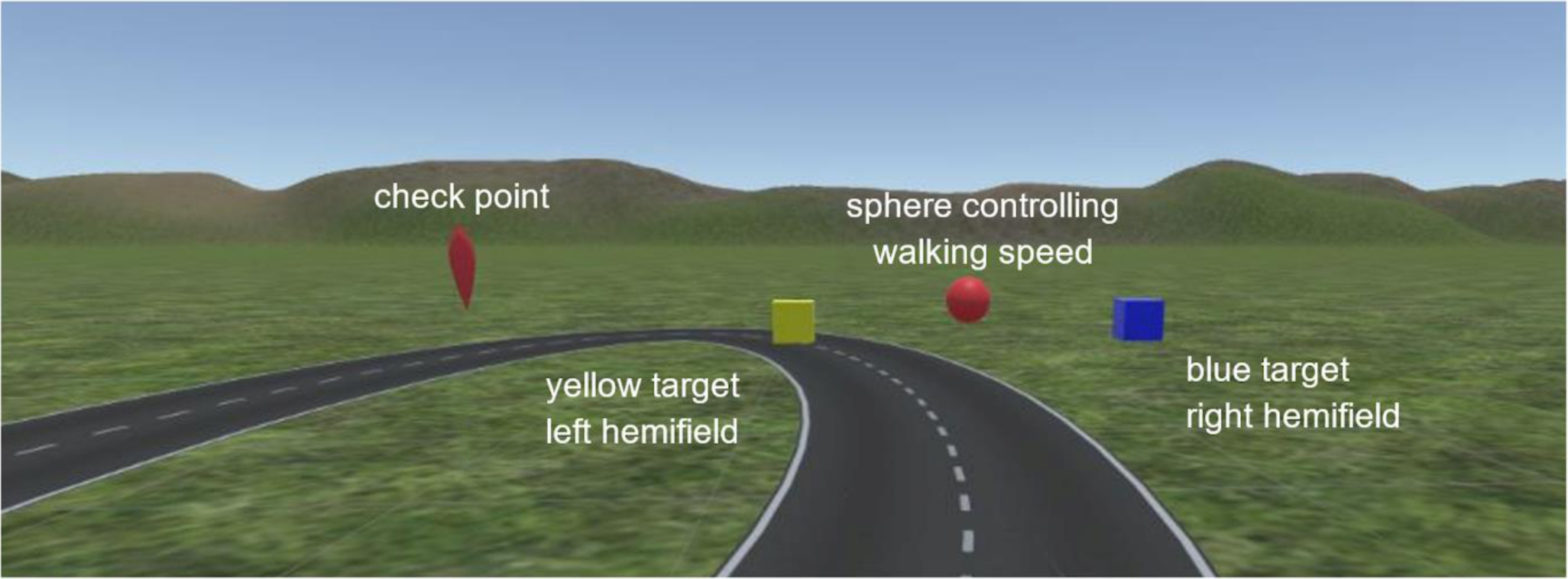
First person view of the participant. A red sphere is placed centrally on the path controlling the walking pace. Next to it, two examples of targets for the visual discrimination task are shown: in the Left hemifield the Yellow target, in the Right hemifield the Blue target. Targets could appear in a randomized fashion in the left and right hemifield and at 15° or 35° of eccentricity at the same height of the participant. On the left, an illustration of the check point that participants had to reach for starting the experiment.

In the main experiment participants performed a visual discrimination task while walking or standing with the movement conditions alternating in six blocks. Each block consisted of 240 trials amounting to 720 trials per movement condition and a total of 1440 trials per participant. The initial movement condition was counterbalanced across participants; fifty percent of participants started in the standing while the other in the walking condition. Between each block, participants were asked to take a break, allowing them to sit down for a few minutes and to flip-up the VR headset. Since the headset display could be flipped up without moving the headset position on the head, the position of electrodes was not affected.

As illustrated in figure 2, a virtual green field was used as the background. The red sphere, controlling for the subject’s walking speed, was placed centrally on the grey path and participants were instructed to keep their gaze towards the sphere at all times to reduce head movement. A yellow cube (*Yellow* condition) or a blue cube (*Blue* condition) were presented for 200 ms in a pseudo-random fashion in the left (*Left* condition) or in the right (*Right* condition) hemifield at 15° or 35° eccentricity (*15°* and *35°* condition respectively). Participants had to press the trigger on the right controller whenever a blue cube was presented and the trigger on the left controller when a yellow cube appeared, creating congruent and incongruent hand-hemifield conditions relatively to the position where the stimuli appeared. If participants responded correctly within a 1.5 second time interval after stimulus presentation, the response was labelled as ‘correct’; vice versa if a wrong trigger was pressed, it was registered as ‘incorrect’; if none of the triggers were pressed, the response was classified as ‘missed’. After each response, a 2000 ms time window was introduced before the presentation of the next stimulus. Task and instructions were identical for both the standing and the walking phase.

Only right-handed participants were recruited, and the walking direction was chosen to be counterclockwise in order to avoid possible effects of handedness on the turning behavior (Angelique et al., 2002; Mohr et al., 2004, 2007; Karim et al., 2016). Second, the height of the sphere and the stimuli were fixed at the height of the participant (right above the eye level) and not to the headset, such that the sphere was not affected by head movement during locomotion. Third, when moving the head away from the red sphere, the sphere stopped moving and changed its color to black until participants turned their heads back in line with the sphere. In this way, we controlled for the head orientation.

### 2.4 EEG recording and preprocessing

EEG recordings were conducted with a 128-channel mobile EEG system (MOVE, Brain Products, Munich, Germany), with a sampling frequency of 1000 Hz. Two EOG electrodes were placed under each eye to measure vertical eye movements and EEG electrodes were placed equidistant according to a custom layout. Impedances of all scalp electrodes were kept below 10Kohm. Raw data was offline processed using MATLAB R2018a (MathWorks, Natick, MA, USA) and EEGLAB 14.1.2b toolbox (SCCN, University of California San Diego, USA, 2018). All preprocessing steps were conducted using the BeMoBIL Preprocessing Pipeline (Klug, 2019) which specifically aims to generalize and simplify the processing of continuous EEG data acquired during MoBI experiments.

The raw EEG data was first filtered to the range of 0.2 Hz and 90 Hz using a finite impulse response (FIR) filter with zero phase (see beemobil_filter()) and resampled to 250 Hz. Subsequently, bad channels were identified and removed by automated rejection using kurtosis (the threshold was set to 5 standard deviations (SDs) from mean kurtosis) and probability functions (with a threshold of 3 SDs from mean probability distribution). Removed channels were interpolated with spherical interpolation and data were subsequently re-referenced to average reference. This “precleaned” dataset was screened for additional spikes and other artifacts (e.g. muscle activity, noise) by visual inspection. Then, for the identification and removal of eye blink artifacts we utilized the procedure of adaptive mixture independent component analysis (AMICA, Palmer et al., 2012; Hsu et al., 2018). To this end, the raw data were bandpass filtered to 1 Hz - 90 Hz to improve the decomposition to independent components (ICs) (Winkler et al., 2015). The resulting decomposition matrices representing the weights and spheres obtained from the AMICA procedure were applied on the “precleaned dataset” (described above) to allow for further component investigation in IC space. ICs representing eye movements (e.g. blinks) were removed based on visual inspection of their component activity profile as well as component power spectra (Makeig et al., 1996; Chaumon et al., 2015). Additionally, prior to IC rejection, we performed additional plausibility testing of our IC selection in terms of its equivalent dipole location to be certain that they did not reflect any brain activity.

After IC removal and back-projection to channel space, the dataset was further highpass filtered with a frequency of 40 Hz. Finally, the cleaned continuous dataset was epoched with onset of each visual stimulus with a pre-stimulus time of −200 ms to 1000 ms after stimulus presentation, and a baseline correction was performed subtracting the −200ms to 0ms pre-stimulus interval from the signal at each channel and trial. During this step, we additionally accounted for a constant temporal delay of 15 ms that were caused by the Brain Vision RDA interface, the WIFI (MOVE) transmission and an additional delay resulting from the Unity software.

### 2.5 Statistical analysis

In the following, the standing condition is referred to as “*Single-task”* condition, and the walking phase is referred to as “*Dual-task”* condition. Besides differences between *Single-* and *Dual-task* (factor: *‘*Task’), we investigated potential lateralization of behavioral and brain responses (factor: ‘Hemifield’) and differences induced by the color of the target (factor: ‘Target’). Moreover, we investigated potential effects induced by the position where stimuli appeared (factor: ‘Eccentricity’). All statistical analyses were conducted using repeated measures ANOVA. When sphericity assumptions were violated in the Mauchly’s test of sphericity (Mauchly, 1940), the p-values were adjusted following the Greenhouse-Geisser correction (Greenhouse & Geisser, 1959). Additionally, we performed post-hoc tests using the Bonferroni method (Bonferroni, 1936) to correct for multiple comparisons. This statistical procedure was applied to analyze all the following dependent measurements.

#### NASA TLX

To investigate the perceived workload, each participant was asked to fill in the NASA TLX (NASA Task Load Index; Hart et al., 1988) questionnaire always after the third and the fourth experimental block, thus after a *Single-Task* or after a *Dual-Task* phase, dependent on the initial starting condition. The total workload assessed by the questionnaire was divided into the six subjective subscales (‘Items’): *mental demand, physical demand, temporal demand, performance, effort* and *frustration*. We used the questionnaire to assess whether the walking activity had an influence on the subjective mental load. Single items were also investigated separately to investigate a potential impact of the movement condition. Therefore, a 2×6 repeated measures ANOVA for factors ‘Task’ and ‘Items’ was computed.

#### Performance

Three 2×2×2×2 repeated measures ANOVAs were calculated to analyze reaction times, percentage of misses and percentage of incorrect responses. For all three dependent variables, we tested the same within factors: ‘Task’ (*Single-* vs *Dual-task*), ‘Eccentricity’ (*15°* vs *35°*), ‘Hemifield’ (*Left* vs *Right*) and ‘Target’ (*Blue* vs *Yellow*). Reaction times were defined as the time between stimulus onset and button press and analyzed only for correct response trials. Accuracy in task performance was operationalized through the number of missed and incorrect response trials over the total number of trials that remained after artifact correction during the EEG pre-processing. This secured identical trials to enter the performance and EEG statistics. Missed trials were defined as missing responses within the 1.5 seconds after the stimulus onset, and incorrect trials as a deviation from required response pattern (wrong button response).

#### ERPs

The P3 component evoked by the visual discrimination task was analyzed for the midline electrodes of the custom layout that were closest to the standard midline locations. These locations are denoted with an apostrophe (*Fz’, Cz’, CPz’, Pz’, POz’, Oz’*). From this, the centro-parietal regions, where the P3 maximum is usually observed, as well as brain activity at more frontal and occipital regions were investigated on their modulatory effect of the P3 in terms of amplitude, onset time (latency) and topographic distribution. To this end, individual maximum positive peak within a time window of 300-600 ms was detected and the mean value of the peak within an 80 ms (40 ms before and after the time of the peak) range around the peak was computed for further statistical analyses. To have a more accurate estimate of the P3 peak amplitude, the same procedure was repeated with different amplitude windows (maximum peak +/- 10ms, 20ms, 30ms, 40ms). For the final analyses, an 80ms range (maximum peak +/- 40ms) was chosen as the results revealed the same direction of effects and the longer time window was more suitable for including a relatively smeared P3 component. P3 latency (ms) and amplitude (µV) means were analyzed in a full-factorial design (2×2×2×2×s6) with ‘Task’ (*Single-* vs *Dual-task*), ‘Eccentricity’ (*15°* vs *35°*), ‘Hemifield’ (*Left* vs *Right*), ‘Target’ (*Blue* vs *Yellow*) and ‘Channel’ (*Fz’, Cz’, CPz’, Pz’, POz’, Oz’*) as repeated measure factors. The effects of the factors ‘Channel’ and ‘Task’ are in the focus of interest and thus explored more thoroughly in results and discussion. However, complete results are reported in the supplementary material.

#### PSDs

For the PSD (Power Spectral Density) analysis, spectral power (µV ^2^/Hz) in 4-8 Hz (theta), 8-10 Hz (lower alpha), 10-12 Hz (upper alpha), 12-30 Hz (beta), and 30-40 Hz (gamma) band were extracted from the same stimulus-locked epochs used for the ERPs analysis and averaged across each condition. Again, here we focused primarily on the central midline electrodes (*Fz’, Cz’, CPz’, Pz’, POz’, Oz’*): a 2×2×2×2×6 repeated-measures ANOVA was computed for each defined frequency band on factors ‘Task’ (*Single-* vs *Dual-task*), ‘Eccentricity’ (*15°* vs *35°*), ‘Hemifield’ (*Left* vs *Right*), ‘Target’ (*Blue* vs *Yellow*) and ‘Channel’ (*Fz’, Cz’, CPz’, Pz’, POz’, Oz’*). Given our hypotheses, the presentation of results focuses only on the effects of the factors ‘Channel’ and ‘Task’ for each of the frequency bands. The complete results are reported in the supplementary material.

## RESULTS

### 3.1 Subjective measures

#### NASA TLX

The 2×6 repeated-measures ANOVA computed on the NASA TLX subscales did not yield any significant main effect for ‘Task’ or ‘Items’. Only the interaction between ‘Task’ and ‘Item’ was statistically significant (F_5,11_ = 2.766, p = .031, η^2^p = .116). The data revealed a tendency for higher ratings regarding the mental, physical and temporal demand in the *Dual-Task* condition while there was a tendency towards higher performance, effort and frustration scores in the *Single-task* situation. However, corrected post-hoc tests did not reveal significant differences between the conditions (*Single-Task* vs *Dual-Task*) for single items.

### 3.2 Performance measures

#### Reaction time

the analysis of reaction time, we found no significant main effect of ‘Task’ on response times. However, a significant main effect for the factor ‘Eccentricity’ was observed (F_1,17_ = 99.3, p < .001, η^2^p = .825) revealing increased reaction times for stimuli appearing at *35*° eccentricity as compared to *15*° eccentricity. Significant effects were observed also for the factors ‘Target’ (F_1,17_ = 8.04, p < .05, η^2^p = .277) and ‘Hemifield (F_1,17_ = 8.77, p < .01, η^2^p = .295), and for their interaction (F_1,17_ = 43.24, p < .001, η^2^p = .673). Post-hoc comparisons for the interaction between ‘Target’ and ‘Hemifield’ showed increased reaction times when responding with the left hand to *Yellow* targets that appeared in the *Right* hemifield as compared to the same targets that appeared in the *Left* hemifield (p < .001), and compared to *Blue* targets that appeared in the *Right* hemifield and that were responded to with the right hand (p < .001). Similarly, the reaction time was longer for responses with the right hand to *Blue* targets when they appeared in the *Left* hemifield as compared to the same targets when appearing in the *Right* hemifield (p < .001), and to *Yellow* targets appearing in the *Left* hemifield (p < .05). Finally, reaction time were about 23ms longer when responding to Yellow targets with the left hand appearing in the Left hemifield compared to responding to Blue targets with the right hand appearing in the Right hemifield (p < .001).

#### Accuracy

The analysis of response accuracy revealed no significant main effect of ‘Task’ on the percentage of incorrect trials or the percentage of missed trials. However, a significant main effect of ‘Eccentricity’ for the number of missed targets (F_1,17_ = 5.94, p = .024, η^2^p = .221) and incorrect responses (F_1,17_ = 8.35, p < .01, η^2^p = .285) was found. A higher percentage of incorrect responses (4.62%) and missed stimuli (1.68%) was observed when stimuli appeared more peripherally as compared to more centrally presented stimuli (respectively, 3.59% of incorrect responses and 0.75% of missed). Moreover, the analysis of incorrect responses demonstrated a significant interaction between the factors ‘Target’ and ‘Hemifield’ (F_1,17_ = 20.52, p < .001, η^2^p = .494). The pattern replicated the same effect that was already observed for response times. Specifically, significant differences were observed when detecting *Blue* targets (with the right hand) in the *Left* hemifield (6.7% of incorrect trials) as compared to detecting the same targets (with the right hand) in the *Right* hemifield (p < .001; 2.77% of incorrect trials) and *Yellow* targets (with the left hand) in the *Left* hemifield (p < .001; 2.66% of incorrect trials). At the same time, detecting *Yellow* targets (with the left hand) in the *Right* hemifield (4.31% of incorrect trials) led to significantly more incorrect trials as compared to detecting the same *Yellow* targets (with the left hand) in the *Left* hemifield (p < .05; 2.66% of incorrect trials) and *Blue* targets (with the right hand) in the *Right* hemifield (p < .01; 2.77% of incorrect trials). Finally, detecting *Yellow* targets with the left hand in the *Right* hemifield yielded higher percentage of incorrect responses (1.36% of incorrect trials) when compared with detecting *Blue* targets with the right hand in the *Left* hemifield (p < .01; 1.11% of incorrect trials).

### 3.3 ERPs (Event-Related Potentials)

#### P3 Latency

The 2×2×2×2×6 ANOVA computed on the P3 latency (ms) evoked by the visual discrimination task yielded a significant main effect only for the factor ‘Channel’ (F_5,8_ = 7.11, p = .003, η^2^p = .253). No significant effect was found neither for the factor ‘Task’ (F_1,8_ = 2.63, p > .1, η^2^p = .112) nor for its interaction with ‘Channel’ (F_5,8_ = 1.4, p > .2, η^2^p = .063).

#### P3 amplitude

Significant main effects for the factors ‘Channel’ (F_5,8_ = 11.07, p < .001, η^2^p = .345), and ‘Task’ (F_1,8_ = 14.16, p < .01, η^2^p = .403) were observed. These main effects were qualified by their interaction (F_5,8_ = 4.053, p < .01, η^2^p = .162). Post-hoc test showed significant differences in P3 amplitude for the channels CPz’ (p < .05), Pz’ (p < .001) and Oz’ (p < .001). A strong P3 amplitude reduction was observed in all posterior electrodes in the *Dual-Task* as compared to the *Single-task* condition (*CPz’ Single-Task*: M = 5.56, SD = 2.29; *Dual-Task*: M = 5.07, SD = 2.46; *Pz’ Single-task*: M = 3.2, SD = 2.41; *Dual-Task*: M = 2.62, SD = 2.64; *Oz’ Single-task*: M = 3.96, SD = 2.31; *Dual-Task*: M = 3.34, SD = 2.13).

In addition, but not in the focus of interest for this study, significant effects were observed also for ‘Target’ (F_1,8_ = 5.54, p < .05, η^2^p = .209), for the interaction ‘Target’ by ‘Hemifield’ (F_1,8_ = 5.04, p < .05, η^2^p = .194) and for the three-way interaction between ‘Target’, ‘Hemifield’ and ‘Task’ (F_1,8_ = 7.6, p < .05, η^2^p = .266). Significant differences in P3 amplitude between congruent and incongruent hand-hemifield response conditions were observed when responding to *Blue* targets in the *Left* hemifield with the right hand as compared to responding to the same targets with the right hand when they appeared in the *Right* hemifield (p < .01) and responding with the left hand to *Yellow* targets that appeared in the *Left* hemifield (p < .001). Similarly, responding to *Yellow* targets that appeared in the *Right* hemifield with the left hand led to significantly lower P3 amplitudes as compared to responding with the left hand to the same *Yellow* targets in the *Left* hemifield (p < .01). As revealed by post-hoc comparisons for the interaction ‘Task’ by ‘Target’ by ‘Hemifield’, the differences in P3 amplitude linked to congruent and incongruent hand-hemifield response conditions were observed only in the *Dual-task*. Within this task condition, a lower P3 amplitude was observed when detecting *Blue* targets in the *Left* hemifield and responding with the right hand as compared to responding to the same targets in the Right hemifield but had to respond with the right hand (p < .001) and responding with the left hand to *Yellow* targets that appeared in the *Left* hemifield (p < .001). Finally, responding with the left hand to *Yellow* targets that appeared in the *Right* hemifield led to significantly lower P3 amplitudes as compared to responding with the left hand to the same *Yellow* targets in the *Left* hemifield (p < .05) and responding with the right hand to *Blue* targets appearing in the *Left* hemifield (p < .01).

### 3.4 PSDs (Power Spectral Densities)

#### Theta

Significant main effects were observed in the theta frequency band (4-8 Hz) for the factors ‘Channel’ (F_5,8_ = 12.79, p < .001, η^2^p = .379) and ‘Task’ (F_1,8_ = 39.98, < .001, η^2^p = .656). In addition, a significant interaction effect between ‘Channel’ and ‘Task’ was observed (F_5,8_ =, p < .01, η^2^p = .140), revealing a significantly lower theta power in the *Single-task* as compared to the *Dual-Task* condition for all the midline channels Fz’ (p < .001), Cz’ (p < .001), *CPz’* (p < .001), *Pz’* (p < .001), *POz’* (p < .001), and *Oz’* (p < .001).

#### Lower alpha

Significant results for the factor ‘Channel’ (F_5,8_ = 9.05, p < .001, η^2^p = .301) and for its interaction with the factor ‘Task’ (F_5,8_ = 4.59, p < .01, η^2^p = .180) were found for the lower alpha frequency range (8-10 Hz). Post-hoc analysis revealed lower alpha power in the *Dual-Task* condition as compared to the *Single-task* condition, reaching significance at channels *CPz’* (p < .05), *Pz’* (p < .05) and POz’ (p < .001) but not Fz’, Cz’ and Oz’.

#### Upper alpha

In the 10-12 Hz range, significant effects were only found for ‘Channel’ (F_5,8_ = 3.62, p < .05, η^2^p = .147). The interaction ‘Channel’ x ‘Task’ also reached significance (F_5,8_ = 4.2, p < .05, η^2^p = .167), revealing a significantly lower average power for the upper alpha band in the *Dual-Task* condition as compared to the *Single-task* condition for electrodes *Fz’* (p < .001), *Cz’* (p < .001), *CPz’* (p < .001) and *Pz’* (p < .05).

#### Beta

PSD differences in the beta frequency band (12-30 Hz) were observed, with significant main effects of interest for the factors ‘Channel’ (F_5,8_ = 10.72, p < .001, η^2^p = .338) and ‘Task’ (F_1,8_= 4.6, p < .05, η^2^p = .180). A significant interaction effect for the same factors also emerged (F_5,8_ = 9.16, p < .001, η2p = .304). Post-hoc tests revealed a significant lower power in the beta band for the *Single-task* as compared to *Dual-Task* for Pz’ (p < .001), POz’ (p < .001) and Oz’ (p < .001), and the opposite trend in CPz’ (p < .001).

#### Gamma

Power in the gamma frequency band (30-40 Hz) yielded significant main effects for the factors ‘Channel’ (F_5,8_ = 14.8, p < .001, η^2^p = .120) and ‘Task’ (F_1,8_ = 21.9, p < .001, η^2^p = .167), and also for their interaction (F_5,8_ = 28.05, p < .001, η^2^p = .205). Post-hoc comparisons revealed a significantly higher gamma power when comparing the *Dual-Task* with the *Single-task* conditions for all midline electrodes: Fz’ (p < .001), Cz’ (p < .001), CPz’ (p < .001), Pz’ (p < .001), POz’ (p < .001), Oz’ (p < .001).

## DISCUSSION

The present study was designed to give further insights into the human brain dynamics of dual-task walking, particularly when the cognitive task taxes the same visual resources that are also required for natural overground walking. To this end, we used a visual discrimination task in VR and analyzed the impact of dual-task walking on cognitive performance and brain dynamics using the MoBI approach. The head mounted VR allowed for a dynamic presentation of stimuli dependent on the participants actual heading in the simulated environment, providing higher ecological validity while, at the same time, ensuring control of potential confounding factors. In this way we were able to address a relevant aspect of mobile cognition which is resource conflicts during walking when visual stimuli in different areas of the visual field are processed.

In this framework, task performance was assessed using reaction times and percentage of incorrect and missed responses as a measure of accuracy. The NASA TLX was performed once after the *Single-task* and once after the *Dual-Task* condition to measure the subjectively perceived mental load. In addition, the P3 component and the PSDs evoked by the cognitive task were analyzed. We hypothesize that CMI is likely to affect response time and accuracy in a visual discrimination task, particularly when visual stimuli are to be detected more peripherally. Moreover, a dual-task cost was expected to impact P3 latencies and amplitudes as well as PSDs in the theta and alpha frequency bands, as well as in higher frequencies, as a consequence of the reallocation of attentional resources related to the dual-task load.

### Subjective measures

Our results did not reveal significant differences in the perceived mental load between the *Single-task* and *Dual-Task* conditions. Despite a significant interaction of the factors ‘Task’ and ‘Item’, no significant difference between the scores for any of the single items were found in the post-hoc contrasts. Therefore, against our hypothesis, the overall results from the NASA TLX reflected a comparable workload for the *Dual-Task* and the *Single-task* conditions. Only a general tendency for higher mental, physical and temporal demand in the *Dual-Task* situation and a tendency towards higher performance, effort and frustration scores in the *Single-task* situation were observed. However, none of the single items reached significance when the appropriate Bonferroni correction was applied. This might have been due to the secondary task, which, arguably, was particularly easy for the young healthy population under investigation. Nonetheless, it is of interest to point out that even for such a low-demanding cognitive task that did not reveal any differences in perceived mental load, the behavioral measures and brain dynamics revealed systematic task-related differences.

### Performance measures

The absence of a main effect of the factor ‘Task’ for all dependent measures of performance likely reflects the relatively low complexity of the chosen secondary cognitive task. Interestingly, 504 overall missed trials were counted in the *Dual-Task* condition, in contrast to 258 missed trials observed in the *Single-task* condition. Even though this is a clear tendency for a behavioral cost induced by the dual-task walking, this trend failed to reach significance.

By contrast, our hypothesis regarding the impact of the target eccentricity was supported. Reaction times were significantly slower when responding to stimuli appearing at *35°* as compared to *15°* eccentricity. In addition, a higher miss rate and higher incorrect responses were observed for stimuli at *35°* eccentricity as compared to those presented at *15°* eccentricity. Previous studies already demonstrated such an eccentricity effect in vision and attention (Carrasco et al., 1995; Staugaard et al., 2016). Some argue that the effect can be explained by the neurophysiological differences between central and peripheral vision (Carrasco & Frieder, 1997). Others state that this could reflect a central bias in the allocation of attentional resources (Wolfe et al., 1998; Brown et al. 2005). Either way, it has long been established that the fovea has higher visual acuity, spatial resolution and contrast sensitivity relatively to the periphery (Carrasco & Frieder, 1997), and it also appears to be favoured in the distribution of attention (Wolfe et al., 1998; Brown et al. 2005). The eccentricity effect was observed in our novel virtual dual-task walking paradigm as well, proving this approach to replicate more traditional laboratory setups during natural overground walking and challenging the recent results reported by Cao and Haendel (2019). As shown by our results, the position of a salient and attention demanding stimulus can impact visual information processing with higher accuracy and faster responses in the central visual field.

An interaction effect between ‘Target’ and ‘Hemifield’ was found both for reaction times and incorrect responses. This can be explained with the Simon effect (Simon, 1969) that predicts increased reaction times and a higher percentage of incorrect responses when targets have to be responded to with an incongruent hand-hemifield response assignment (*Yellow-Right* or *Blue-Left*) as compared to the conditions with congruent hand-hemifield responses (*Yellow-Left* or *Blue-Right*).

### ERPs (Event-Related Potentials)

Even though the performance measures did not provide any direct evidence for a main effect of increased effort in the *Dual-Task* condition, the ERP results clearly indicated differences in neural processing between *Single-* and *Dual-Task*. Indeed, a significant impact of the ‘Task’ condition was observed for the P3 amplitude, showing a strong reduction in the *Dual-Task* as compared to the *Single-task* (figures 4 and 5). These effects differed topographically and were most pronounced over posterior leads (CPz’, Pz’, Oz’). This P3 amplitude reduction with a posterior maximum replicated previous results on P3 amplitude reductions in dual-task walking scenarios (De Sanctis et al., 2014; Malcolm et al., 2015; Ladouce et al., 2019; Reiser et al., 2019). However, in contrast to the above mentioned papers, the P3 onset latency did not differ between *Single-* and *Dual-task* conditions.

**Fig. 3:**
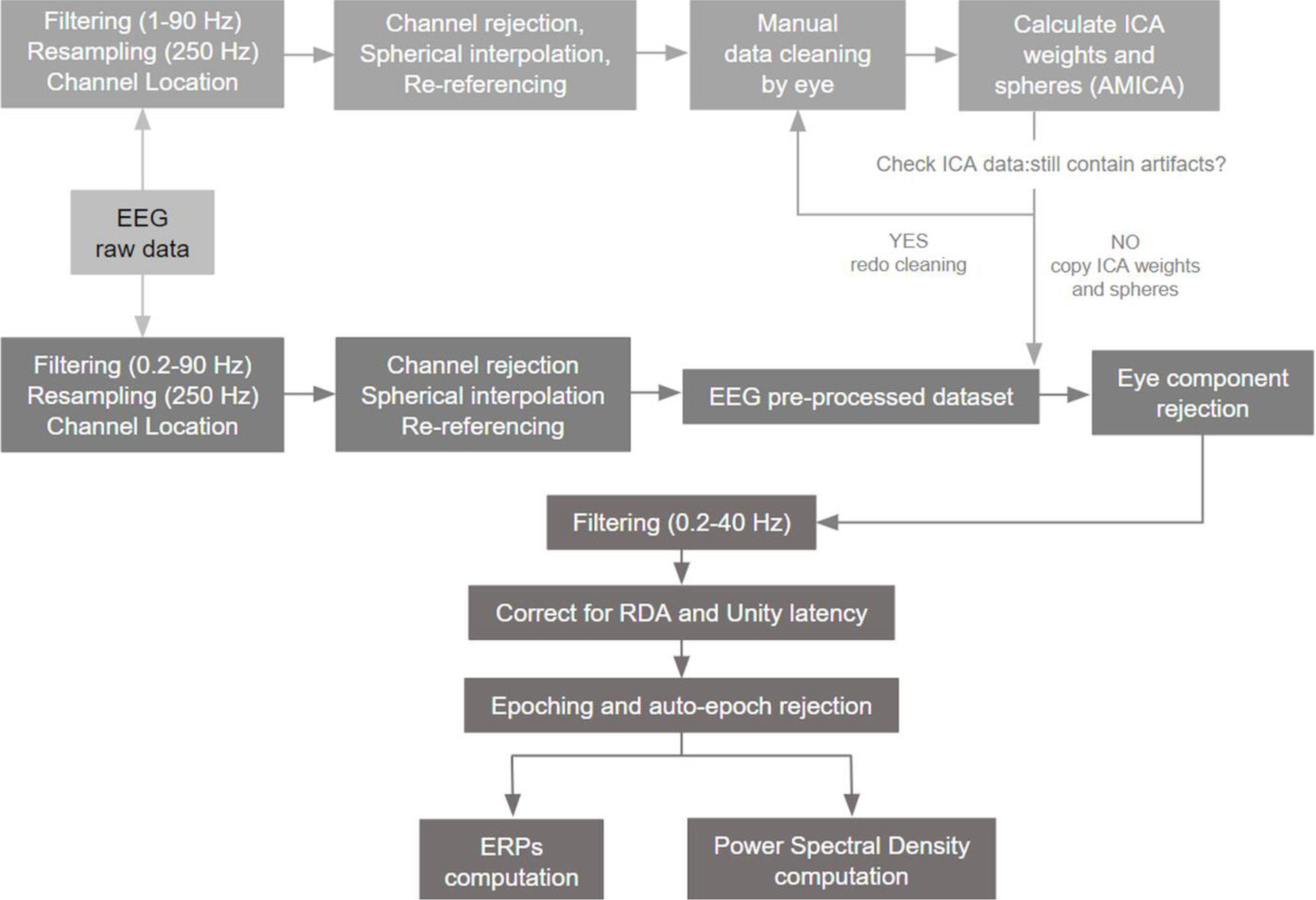
Overview of the preprocessing pipeline for the continuous raw EEG data.

**Fig. 4:**
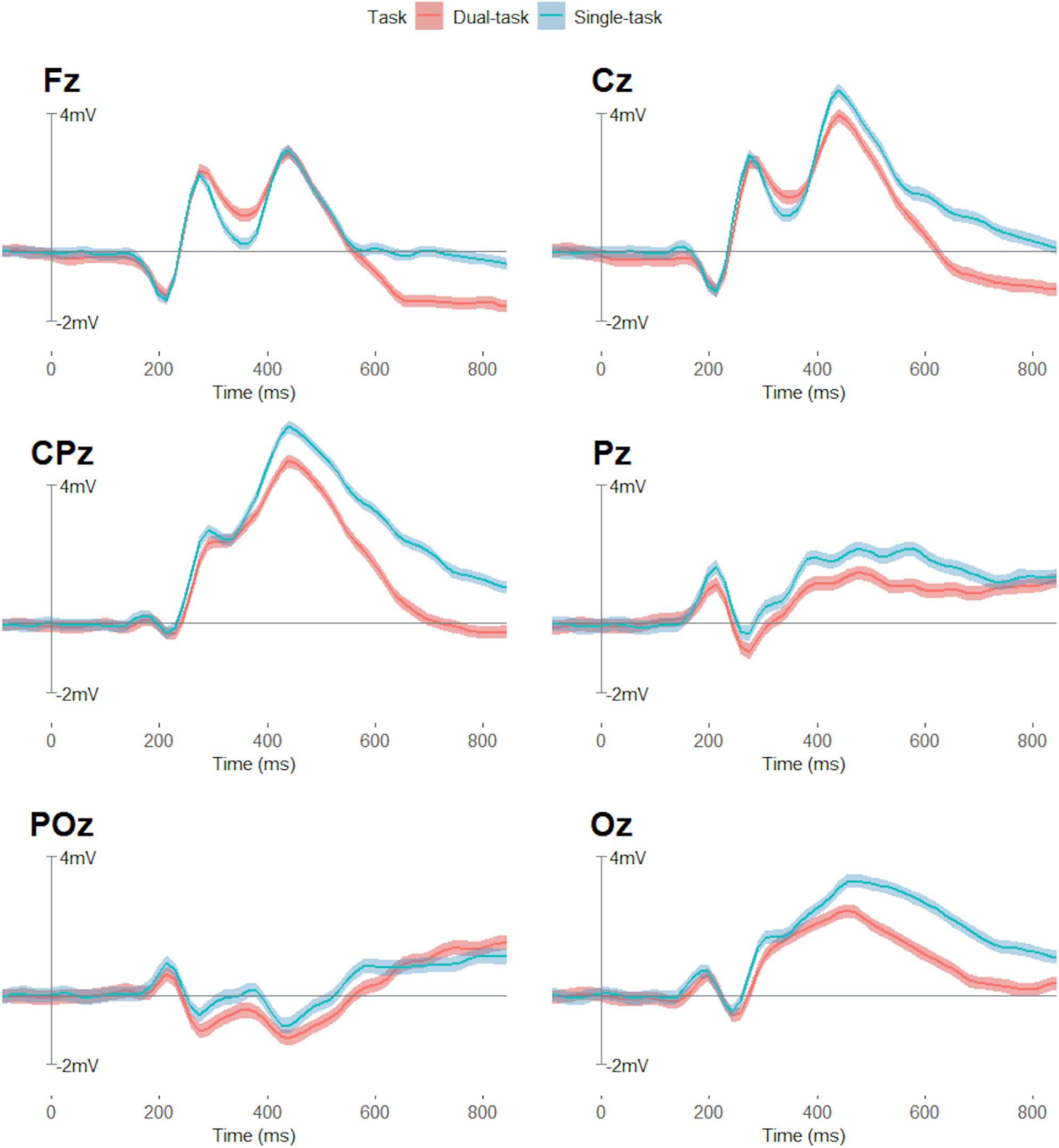
ERP components extracted for each midline channel. Blue curve depicts the ERP waveform for correct responses during the *Single-task* condition, while the red curve during the *Dual-task* condition. The shadow around each component reflects the relative standard deviation.

**Fig. 5:**
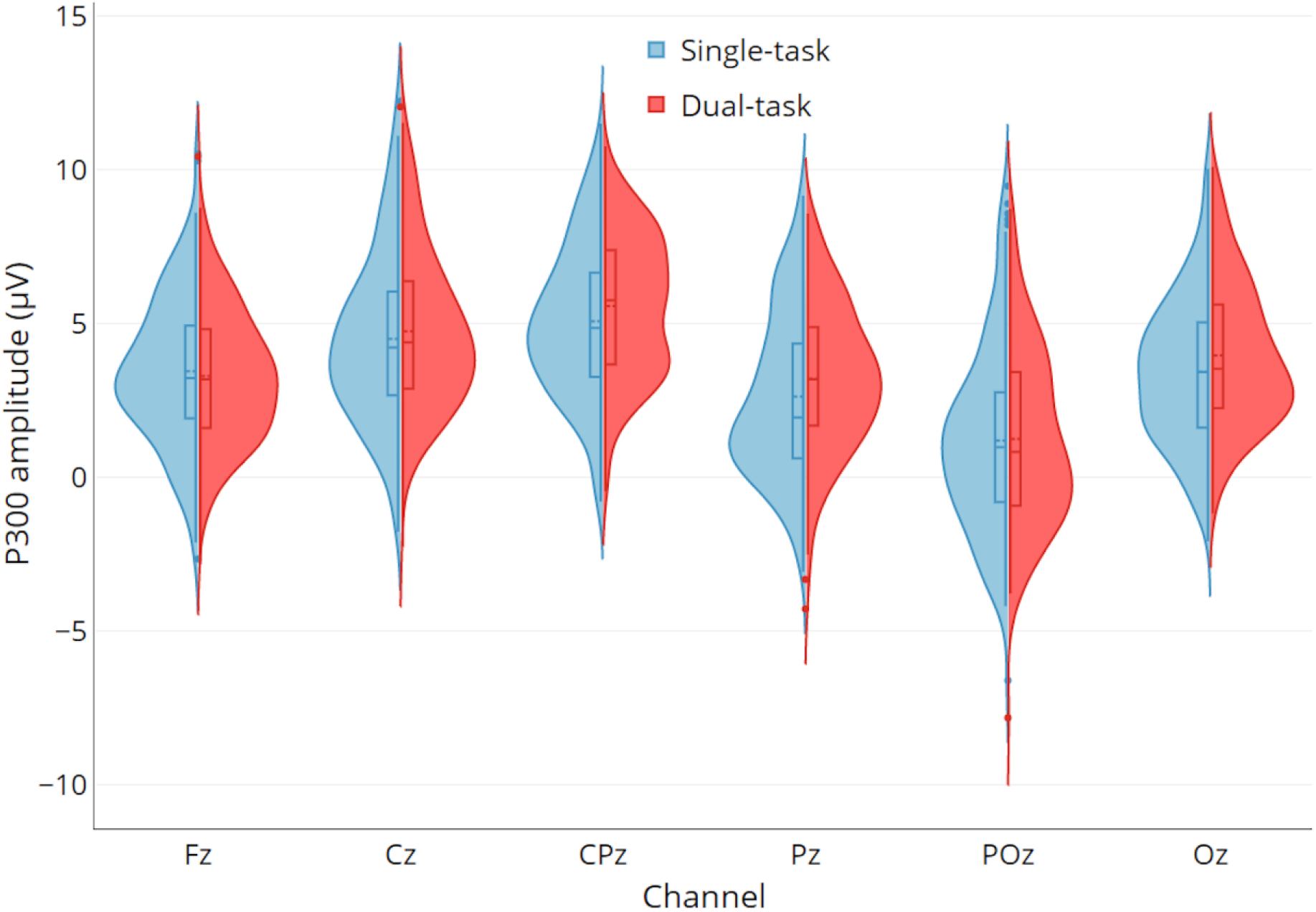
P300 amplitude distributions evoked by the visual discrimination task. Violin plots are relative to the ‘Task’ condition and separated by ‘Channel’ (on the X axis).

Furthermore, it is important to note that the EEG recordings at POz’ were particularly noisy and thus difficult to interpret. As depicted in figure 7, the straps of the Mixed Reality goggles were situated precisely over this electrode, likely inducing significant pressure and mechanical noise. This was confirmed by the P3 signal-to-noise ratio (SNR) which was calculated in each of the midline channels dividing the ERP amplitude by the standard deviation in the prestimulus interval (Debener et al., 2008). For POz’, the SNR was significantly lower compared to the SNR at all other midline electrodes except for channel Cz’ (POz’-Fz’: p < .05; POz’-Cz’: p = .27; POz’-CPz’: p < .001; POz’-Pz’: p < .001; POz’-Oz’: p < .01). It is thus reasonable to assume that the ERPs at POz’ were compromised by the constant mechanical pressure of the VR headset, making it difficult to detect reliable peak onset-latencies.

**Fig. 6:**
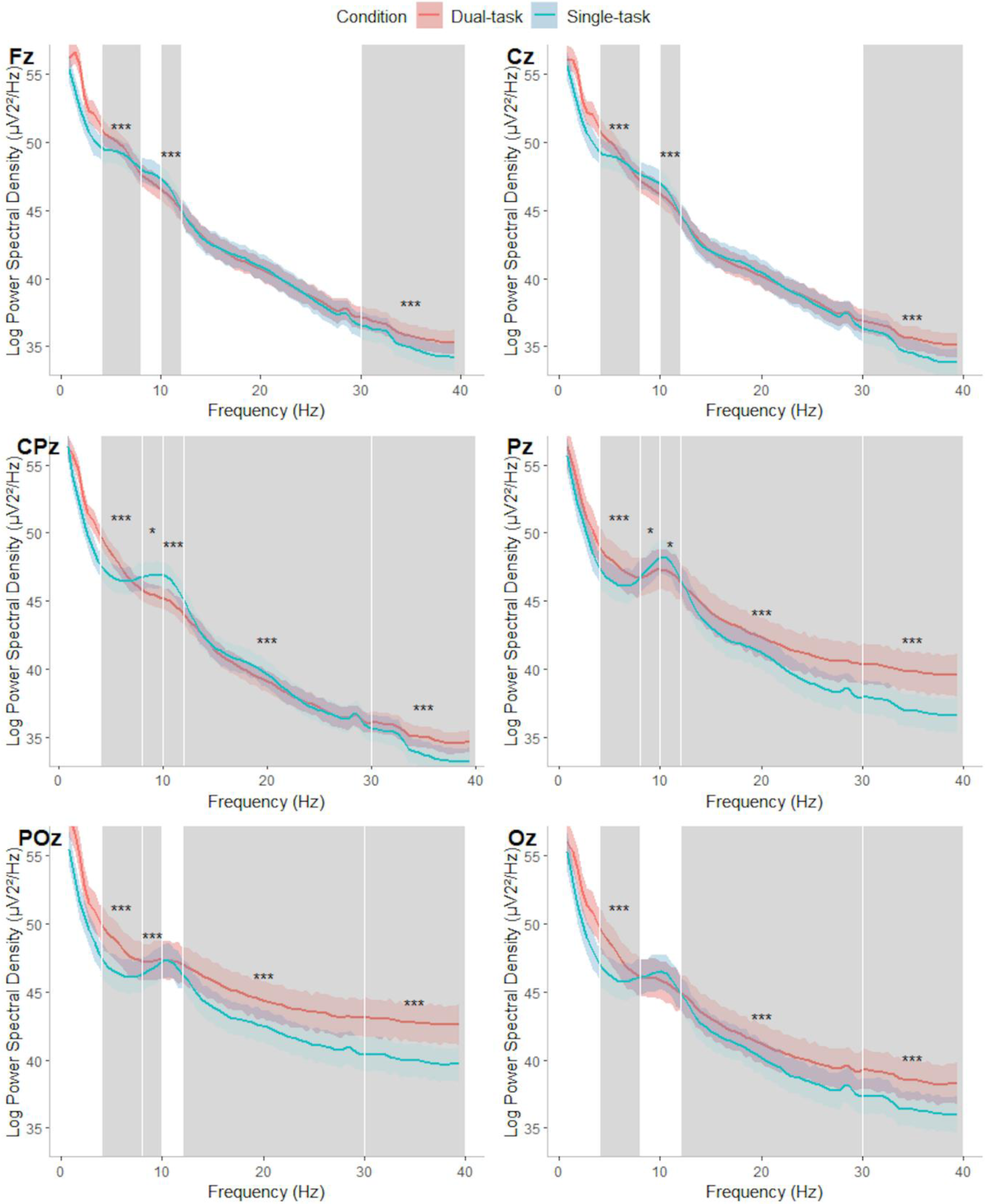
Power Spectral Densities (PSDs) extracted from EEGLAB for each of the midline channels are depicted relatively to the ‘Task’ condition. All the spectra were divided in five bands (theta: 4-8 Hz; lower alpha: 8-10 Hz, upper alpha: 10-12 Hz; beta: 12-30 Hz; gamma: 30-40 Hz), independently analyzed through five 2×2×2×2×6 repeated measures ANOVAs. Bands highlighted in grey have yielded significant effects and are complemented by stars indicating the significance level of the test (*: p ≤ 0.05; **: p ≤ 0.01; ***: p ≤ 0.001).

**Fig. 7:**
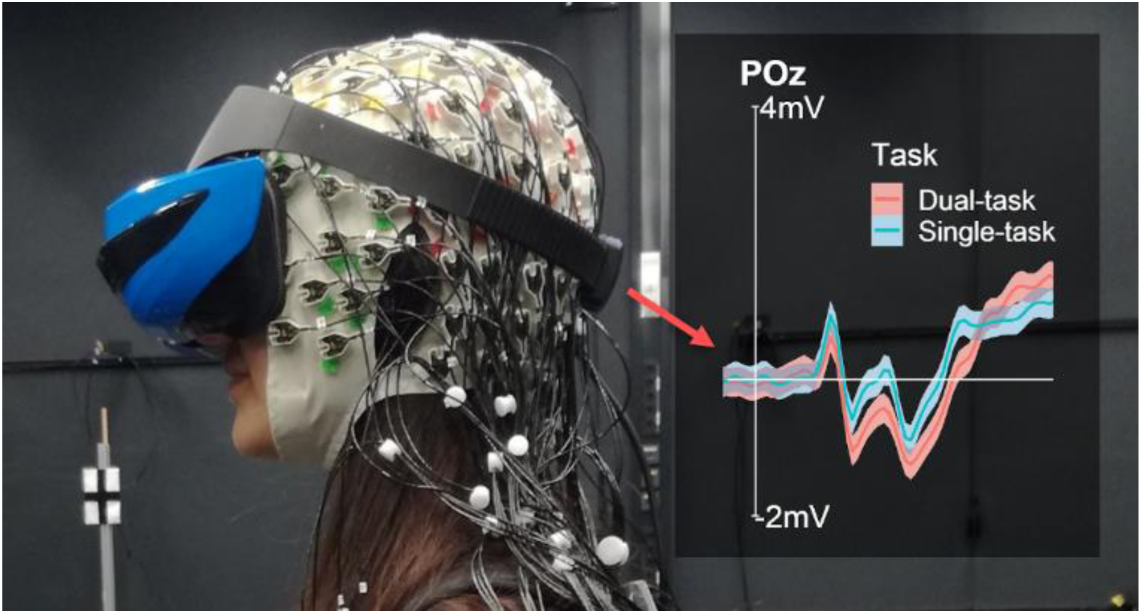
Depiction of the EEG waveform of the POz electrode that is contaminated by noise caused by the pressure of the ASUS Mixed Reality headset

Finally, the same Simon effect (Simon, 1969) found in the reaction times and the response accuracy was observed for the P3-component as well. A smaller P3 amplitude was observed for incongruent hand-hemifield responses (*Blue Left* and *Yellow Right*) in contrast to congruent responses (*Blue Right* and *Yellow Left*). These results have repeatedly been observed in previous static setups (Ragot et al., 1990; Zhou et al., 2004; Melara et al., 2008) and are here replicated for a naturalistic walking task. Interestingly, as shown by the three-way interaction between ‘Task’, ‘Target’ and ‘Hemifield’, the incongruence of hand-hemifield response did not affect P3 amplitude within the *Single-task* but only within the *Dual-task* condition. These results indicate a greater difficulty for the *Dual-task* walking condition when participants needed to respond to targets with an incongruent hand-hemifield assignment. An increased difficulty and associated P3 amplitude in the walking as compared to the standing condition can be explained by a resource conflict. Due to increased resources necessary for walking, the additional button press response created stronger interference compared to the standing (Ruffieux et al., 2015). Such a resource conflict is reduced for standing in which the Simon effect was thus marginalized. While the Simon effect in our setup is noteworthy, the more important finding was an impact of the hand-hemifield incongruence on brain dynamics only during natural overground walking but not while standing.

### PSDs (Power Spectral Densities)

Gradually proceeding from lower to higher frequency bands, the present study replicated an inverse modulation of **theta** and **alpha** activity related to the task demand. An increased theta power in the *Dual-* as compared to the *Single-task* was observed over frontal and centro-parietal areas spreading out to occipital sites. Moreover, a decreased alpha power has been observed as well, which was prominent from central sites up to parieto-occipital regions for lower alpha frequencies, and from frontal up to parietal regions for the upper alpha frequency range. These modulations have previously been observed in a dual-task walking paradigm of Beurskens et al. (2016) and were replicated in the present study during walking overground in a virtual scenario, providing evidence for higher demands during dual-task walking reflected in lower alpha activity in frontal and central brain regions. In contrast to the study by Beurskens and colleagues (2016), the observed alpha decrease was not limited to frontal and central sites but was observed over widespread regions from frontal to parieto-occipital midline channels.

Moving towards higher frequencies, we further observed an increase of **beta** power particularly pronounced over parietal and occipital leads. Previous static paradigms described a link between beta activity and attentional modulation in the visual system (Belyavin & Wright in 1987; Gola et al., 2013), and also a relation with increasing effort to stay alert (Lafrance et al., 2000; Boksem et al., 2005). Less consistent patterns for beta modulation were observed in paradigms that involved walking participants. Here, beta power has been observed to increase (Presacco et al., 2011) and sometimes to decrease (Pizzamiglio et al., 2017). Moreover, while walking, a more pronounced increase of beta power has been observed for a secondary motor task condition than an additional secondary cognitive task (Beurskens et al., 2016). Given the highly contradictory observations reported in several MoBI dual-task walking paradigms, it is difficult to dissociate possible beta modulations specifically related to the cognitive or motor demand. In addition, considering that a walking phase without additional cognitive task was not included in our experimental design, no clear conclusion can be drawn from our results. Future studies will have to systematically address the impact of different dual task modalities and response requirements on beta modulations during walking.

Finally, we observed a strong increase of **gamma** activity when walking for all midline electrodes. This power increase became increasingly pronounced from frontal to occipital sites. Gamma has previously been shown to be related to body instability, which can occur in dynamic situations like walking (Slobounov et al., 2009). Increased gamma activity was also observed when walking and performing an additional cognitive task (Marcar et al., 2014). Both factors were involved in the present study replicating previous results and pointing to a role of gamma activity in the stabilization of posture as well as additional tasks during walking. A functional role of gamma in CMI during natural overground walking however, cannot be drawn from the present study as we did not systematically control between these two factors. In addition, interpretations of gamma activity have to consider a potential confound due to (neck) muscle activity, which is likely to contaminate the surface EEG signal when walking (Whitham et al., 2008). Since gamma modulations are usually related to the motor activity itself, future studies will have to replicate CMI effects including different conditions of motor and cognitive tasks and how these impact gamma activity.

### Principal contributions, limitations and future directions

The fundamental role of vision during motion has been stated clearly (Imai et al., 2001; Nomura et al., 2005; Marigold & Patla, 2008): when walking, visual information processing is required to secure a stable gait pattern. However, due to fixed desktop restrictions, most previous investigations marginalized the possible impact of visual information processing during walking on a treadmill. As an option for studying the neural mechanisms underlying CMI during walking, MoBI is certainly a suitable method (Gramann et al., 2010; Presacco et al., 2011; Debener et al., 2012; De Sanctis et al., 2014; Marcar et al., 2014; Malcolm et al., 2015; Beurskens et al., 2016; Pizzamiglio et al., 2017; Ladouce et al., 2019; Reiser et al., 2019) confirmed by the results of the presented study. In addition, the incorporation of a VR system to MoBI setups opens up a wide range of possibilities for studying visual dual-task walking in all its facets.

Within the present virtual framework, the main question was whether and how attention is allocated when discriminating visual stimuli at different eccentricities while standing and walking. In this context, the perceived mental load during the *Single-* and the *Dual-task* did not differ, and neither did the performance. From a neural perspective, instead, our results demonstrated that simply walking overground at a natural speed already interferes with the execution of a low-demanding cognitive task in the visual domain, even in the absence of performance cost. This was revealed by a P3 amplitude reduction when executing the cognitive task in motion as compared to standing. This was also reflected in the frequency domain with increasing theta and decreasing alpha power over widespread regions of the brain. These mechanisms are known to capture attentional variations accompanying task demand variations (Klimesh et al., 1999; Polich, 2007) and were here registered during more natural cognitive processing while walking. Even though important modulations at higher frequencies were also observed, whether and how those are related to cognitive and motor load is still under debate (Belyavin & Wright, 1987; Gwin et al., 2011; Marcar et al., 2014).

Thus, on the same interpretative line of previous works on mobile cognition (Debener et al., 2012; De Sanctis et al., 2014; Malcolm et al., 2015; Beurskens et al., 2016; Pizzamiglio et al., 2017; Ladouce et al., 2019; Reiser et al., 2019), our results reflect CMI to take place during *Dual-task* walking even when involving only a low-effort visual discrimination task. As an addition to previous works, it was also demonstrated that the above-mentioned neural markers can be used to identify changes in attention during active behaviors involving visual processing in VR during overground walking. However, the visual discrimination task used in the present study was particularly easy for the young population which did not show performance costs related to the Task condition. Future investigations implementing an improved virtual design with more challenging cognitive and motor tasks will have to investigate the roles of both cognition and motion in brain dynamic modulations, still controlling relevant experimental features in an ecologically valid way.

Overall, humans are not static agents passively perceiving changing stimuli from the environment. Every healthy individual has the power to actively move in the surroundings, processing environmental information, preparing actions, and interacting with the external world. This complex intersection between environmental and bodily dynamics needs to be emphasized in experimental contexts rather than being marginalized. The combination of the MoBI with VR, while still not representing cognitive processes in the real world, allows for more ecologically valid dual-task walking investigations, taking a step toward the investigation of more natural and active behaviors involving visual processing. It allows for simulating close-to-reality situations which have a moderate degree of conformity with the real world, but allow for investigating even potentially dangerous contexts. For instance, crossing the street while texting is an undemanding but possibly unsafe behavior which can be safely simulated in VR. This kind of setup allows for enhancement of ecological validity when studying natural behaviors in real world contexts, particularly those involving visual attention while moving in the surrounding. We are confident that the implementation of head mounted VR systems in dual-task scenario will provide an important contribution to better understand ‘natural cognition’, representing the step from the laboratory setting to the real world.

## ACKNOWLEDGEMENTS

Sincere thanks go to Marius Klug, Lukas Gehrke and Zakaria Djebbara for their technical support on MoBI systems. We also thank Zakaria Djebbara for further assistance in programming the virtual environment and during data collection. Finally, we thank Prof. Marco Zorzi for his guidance and additional endorsement. This research was supported by the BeMoBIL (Berlin Mobile Brain/Body Imaging Lab) and was carried out within the scope of the project “use-inspired basic research”, for which the Department of General Psychology of the University of Padova has been recognized as “Dipartimento di eccellenza” by the Ministry of University and Research.

## AUTHOR CONTRIBUTIONS

KG, CTD and FN were responsible for the early study concept and design. CTD and FN collaborated for programming the virtual environment in Unity. FN was responsible for participant recruitment and data collection. JP, KG and FN gave important contributions to the preprocessing and analysis of EEG data, and all authors contributed to data interpretation. FN initially drafted the manuscript and KG, CTD and JP subsequently provided extensive critical revisions and intellectual insertions. The final version of the publication was approved by all authors.

## DATA ACCESSIBILITY STATEMENT

Upon acceptance of the paper, data and code will be made available on an open-source repository.

## ABBREVIATIONS

CMI: cognitive-motor interference;
ERP: event related potential;
MoBI: mobile brain/body imaging;
VR: virtual reality;
PSD: power spectral density;
SNR: signal to noise ratio

