## Supplemental Table for "Alteration of brain dynamics during natural dual-task walking"

### SUPPLEMENTARY MATERIAL

**Table 1: ANOVA – P300 latency and amplitude**
**Task x Eccentricity x Target x Hemifield x Channel (2x2x2x2x6)**

| | P3 latency | P3 amplitude |
|---|---|---|
| Channel (Fz, Cz, CPz, Pz, POz, Oz) | .000 *** | .000 *** |
| Task | .000 *** | .001 ** |
| Task x Channel | .280 | .009 ** |
| Eccentricity | .000 *** | .255 |
| Eccentricity x Channel | .116 | .051 |
| Hemifield | .000 *** | .677 |
| Hemifield x Channel | .201 | .164 |
| Target | .000 *** | .028 * |
| Target x Channel | .613 | .034 * |
| Task x Eccentricity | .000 *** | .056 |
| Task x Eccentricity x Channel | .186 | .188 |
| Target x Eccentricity | .000 *** | .313 |
| Target x Eccentricity x Channel | .623 | .084 |
| Hemifield x Eccentricity | .000 *** | .926 |
| Hemifield x Eccentricity x Channel | .659 | .377 |
| Target x Hemifield | .000 *** | .036 * |
| Target x Hemifield x Channel | .139 | .455 |
| Task x Target | .000 *** | .079 |
| Task x Target x Channel | .092 | .542 |
| Task x Hemifield | .000 | .219 |
| Task x Hemifield x Channel | .834 | .066 |
| Task x Target x Eccentricity | .000 *** | .409 |
| Task x Target x Eccentricity x Channel | .280 | .157 |
| Target x Hemifield x Task | .000 *** | .012 |
| Target x Hemifield x Task x Channel | .251 | .336 |
| Eccentricity x Hemifield x Task | .000 *** | .044 * |
| Eccentricity x Hemifield x Task x Channel | .264 | .028 * |
| Target x Hemifield x Eccentricity | .000 *** | .055 |
| Target x Hemifield x Eccentricity x Channel | .063 | .538 |
| Task x Target x Hemifield x Eccentricity | .000 *** | .910 |
| Task x Target x Hemifield x Eccentricity x Channel | .078 | .335 |

Significant effects are complemented by stars indicating the significance level of the test
(*: $p \leq 0.05$; **: $p \leq 0.01$; ***: $p \leq 0.001$; ****: $p \leq 0.0001$). P-values are Greenhouse-Geisser corrected.

| Table 2: ANOVA - Power Spectral Densities (PSDs)<br>Task x Eccentricity x Target x Hemifield x Channel (2x2x2x2x6) | | | | | |
|---|---|---|---|---|---|
| | Theta | Lower Alpha | Upper Alpha | Beta | Gamma |
| | 4-8 Hz | 8-10 Hz | 10-12 Hz | 12-30 Hz | 30-40 Hz |
| Channel (Fz, Cz, CPz, Pz, POz, Oz) | .000 *** | .000 *** | .005 ** | .000 *** | .000 *** |
| Task | .000 *** | .400 | .090 | .044 * | .000 *** |
| Task x Channel | .017 * | .011 * | .014 | .000 *** | .000 *** |
| Eccentricity | .000 *** | .917 | .041 * | .079 | .000 *** |
| Eccentricity x Channel | .016 * | .016 * | .059 | .000 *** | .000 *** |
| Hemifield | .000 *** | .173 | .015 * | .148 | .012 * |
| Hemifield x Channel | .000 *** | .000 *** | .017 * | .000 *** | .000 *** |
| Target | .000 *** | .223 | .156 | .069 | .000 *** |
| Target x Channel | .599 | .026 * | .075 | .000 *** | .000 *** |
| Task x Hemifield | .281 | .365 | .912 | .309 | .573 |
| Task x Hemifield x Channel | .002 ** | .054 | .119 | .649 | .002 ** |
| Task x Eccentricity | .564 | .636 | .147 | .011 * | .258 |
| Task x Eccentricity x Channel | .283 | .359 | .081 | .292 | .465 |
| Task x Target | .170 | .059 | .479 | .233 | .070 |
| Task x Target x Channel | .042 * | .819 | .837 | .213 | .166 |
| Target x Hemifield | .326 | .269 | .110 | .607 | .039 * |
| Target x Hemifield x Channel | .041 * | .026 * | .551 | .255 | .018 * |
| Target x Eccentricity | .418 | .080 | .631 | .858 | .012 * |
| Target x Eccentricity x Channel | .855 | .405 | .164 | .948 | .104 |
| Hemifield x Eccentricity | .710 | .836 | .192 | .932 | .652 |
| Hemifield x Eccentricity x Channel | .786 | .437 | .384 | .303 | .457 |
| Target x Hemifield x Task | .000 *** | .872 | .007 ** | .068 | .002 ** |
| Target x Hemifield x Task x Channel | .007 ** | .006 ** | .011 * | .001 ** | .000 *** |
| Task x Target x Eccentricity | .000 *** | .259 | .038 * | .056 | .000 *** |
| Task x Target x Eccentricity x Channel | .137 | .052 | .028 * | .001 ** | .000 *** |
| Task x Hemifield x Eccentricity | .000 *** | .543 | .093 | .016 * | .000 *** |
| Task x Hemifield x Eccentricity x Channel | .165 | .054 | .023 * | .000 *** | .000 *** |
| Target x Hemifield x Eccentricity | .000 *** | .935 | .121 | .077 | .000 *** |
| Target x Hemifield x Eccentricity x Channel | .044 * | .002 ** | .010 * | .000 *** | .000 *** |
| Task x Target x Hemifield x Eccentricity | .005 ** | .334 | .965 | .407 | .158 |
| Task x Target x Hemifield x Eccentricity x Channel | .013 * | .153 | .206 | .427 | .003 ** |

Significant effects are complemented by stars indicating the significance level of the test
(*: $p \leq 0.05$; **: $p \leq 0.01$; ***: $p \leq 0.001$; ****: $p \leq 0.0001$). P-values are Greenhouse-Geisser corrected.

For an interactive DEMO of P3 amplitude data and distribution:
https://chart-studio.plot.ly/~federica.nenna/1/#/plot
